# Sexual dimorphism in the meiotic requirement for PRDM9: a mammalian evolutionary safeguard

**DOI:** 10.1101/2020.03.10.985358

**Authors:** Natalie R Powers, Beth L Dumont, Chihiro Emori, Raman Akinyanju Lawal, Catherine Brunton, Ken Paigen, Mary Ann Handel, Ewelina Bolcun-Filas, Petko M Petkov, Tanmoy Bhattacharyya

## Abstract

In many mammals, genomic sites for recombination are determined by histone methyltransferase PRMD9. Mice lacking PRDM9 are infertile, but instances of fertility or semi-fertility in the absence of PRDM9 have been reported in mice, canines and a human female. Such findings raise the question of how the loss of PRDM9 is circumvented to maintain reproductive fitness. We show that genetic background and sex-specific modifiers can obviate the requirement for PRDM9 in mice. Specifically, the meiotic DNA damage checkpoint protein CHK2 acts as a modifier allowing female-specific fertility in the absence of PRDM9. We also report that in the absence of PRDM9, a PRDM9-independent recombination system is compatible with female meiosis and fertility, suggesting sex-specific regulation of meiotic recombination, a finding with implications for speciation.

**One Sentence Summary:** Sex-specific modulation of a meiotic DNA damage checkpoint limits the requirement for PRDM9 in mammalian fertility.

## Main Text

Meiotic recombination generates genetic diversity and ensures the accuracy of chromosome transmission to the next generation. In many organisms, recombination is not random but occurs preferentially at sites in the genome known as recombination hotspots (1). In a subset of mammals, including mice and humans, the positions of hotspots are determined by the specialized histone methyltransferase PRDM9—a meiosis-specific, DNA-binding zinc finger protein that uniquely trimethylates both lysines 4 and 36 of histone H3 (2-6). This H3K4me3/H3K36me3 double-positive signature is thought to preferentially facilitate recombination at these sites, to the exclusion of other functional elements (1, 6, 7). In the absence of PRDM9, meiotic double-strand breaks (DSBs) occur in normal numbers, but they localize to functional elements enriched for H3K4me3, such as promoters (7). In *Prdm9*-deficient C57BL/6 mice (henceforth B6.*Prdm9*^-/-^), repair of these ectopic DSBs is impaired, leading to prophase I meiotic arrest of both male and female germ cells, with consequent infertility due to failure to produce mature gametes (8). In human males, several point mutations in PRDM9 are significantly associated with non-obstructive azoospermia (9, 10). Together, these observations suggest that PRDM9-dependent recombination is required for successful reproduction.

However, intriguing exceptions have been reported in both mice and a human female. *Prdm9*-deficient PWD/PhJ (henceforth PWD.*Prdm9*^*-/-*^) male mice have normal meiotic prophase I, although they are infertile due to low sperm number (PWD.*Prdm9*^*-/-*^ females are also infertile) (11). Further, some *Prdm9*^*-/-*^ male mice with mixed genetic backgrounds are fertile, suggesting background-dependent genetic modifiers of the phenotype (11). A recent report of a single fertile PRDM9-null woman shows that PRDM9 can be dispensable for fertility in human females (12). Although *Prdm9* is a pseudogene in the canine lineage, both male and female canids reproduce successfully (13, 14). Together, these findings imply that the requirement for PRDM9 in mammalian recombination is more complex than was previously imagined. These findings also suggest that a *Prdm9*-independent recombination initiation pathway must exist even in species using PRDM9; elucidating its mechanisms will help decipher the genetic complexity that is likely involved. In male mice, recombination hotspots on autosomes are PRDM9-dependent, but the recombination hotspots within the pseudoautosomal region of the sex chromosomes are activated by a PRDM9-independent recombination initiation pathway (7). This suggests that both PRDM9-dependent and -independent recombination pathways are active in tandem during mammalian meiosis and might fulfill functions necessary to bypass reproductive bottlenecks due to dysfunctional or absent PRDM9. Although a PRDM9-independent recombination mechanism exists in male mice, it has been unknown if a similar pathway functions in females. In this study, we investigated oogenesis and the effect of genetic background to determine (1) whether there is a sex-specific requirement for PRDM9-dependent hotspot activation, and (2) the possible mechanisms allowing organisms to circumvent the loss of PRDM9 and maintain reproductive fitness.

## Results

### Sex-specific requirement for PRDM9-dependent histone methyltransferase activity in mice

We used CRISPR/Cas9 gene editing to create a point mutation in the PR/SET domain of *Prdm9* (Glu365Pro, henceforth *Prdm9*^*EP*^) in C57BL/6J (B6) mice (Fig. S1A-B). This mutation did not entirely abolish the methyltransferase activity of *Prdm9 in vivo* but was a severe hypomorph, as evident from the ChIP-seq data for H3K4me3 in homozygous mutant spermatocytes (Fig. 1A-E). There was a 7.2-fold reduction in the number of detectable peaks in spermatocytes from homozygous *Prdm9*^*EP*^ males (henceforth B6.*Prdm9*^*EP/EP*^) (Fig. 1B, Table S1), compared to B6 spermatocytes. The intensity of detectable PRDM9-dependent peaks was also sharply reduced; B6.*Prdm9*^*EP/EP*^ showed a 6.2-fold reduction in the mean number of reads per peak relative to B6 (Table S2). Indeed, these peaks were so low that they did not register in aggregation plots of H3K4me3 enrichment at all known PRDM9-dependent sites (Fig. 1C-E), confirming a significant alteration in PRDM9-dependent methyltransferase activity in B6.*Prdm9*^*EP/EP*^ spermatocytes.

**Fig. 1.**
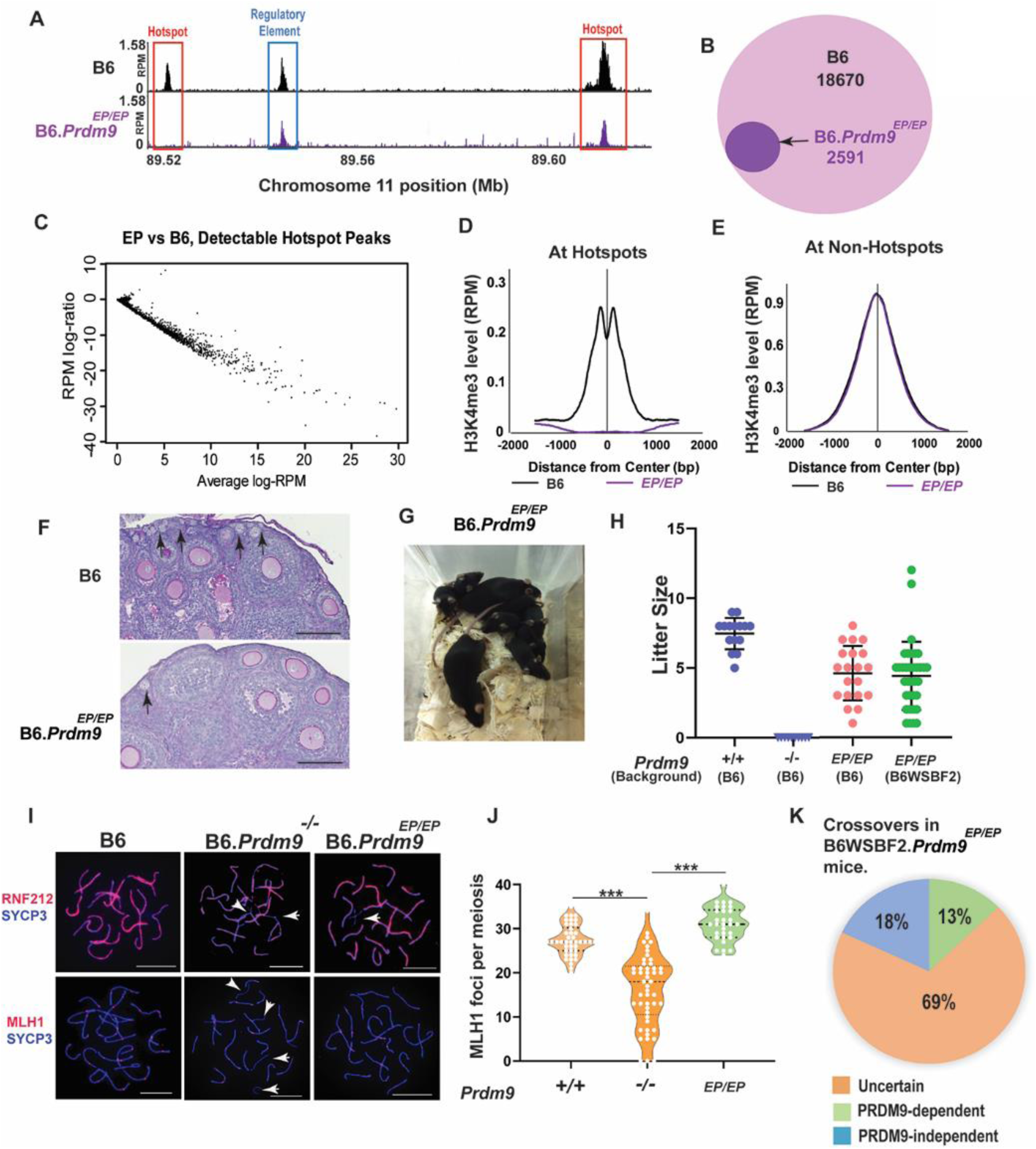
*Prdm9*^EP/EP^ females in the C57BL/6J genetic background are fertile. (A) Genome browser snapshot of H3K4me3 ChIP-seq peaks in wild-type and *Prdm9*^EP/EP^ spermatocytes. One strong and one weaker PRDM9-dependent hotspot are shown, together with one PRDM9-independent peak. (B) Venn diagram showing the number of recombination hotspots with a detectable H3K4me3 peak in wild-type (B6) and *Prdm9*^EP/EP^ spermatocytes. (C) MA plot comparing *Prdm9*^EP/EP^ vs. wild-type signal at PRDM9-dependent H3K4me3 peaks that were detectable in *Prdm9*^EP/EP^ spermatocytes (n = 2,591). (D) Aggregation plot showing normalized average signal intensity (reads per million, RPM) at known PRDM9-dependent H3K4me3 ChIP-seq peaks (n = 18,838) in wild-type and *Prdm9*^EP/EP^ spermatocytes. (E) Aggregation plot showing normalized average signal intensity (reads per million, RPM) at non-hotspot H3K4me3 ChIP-seq peaks (n = 56,030) in wild-type, and *Prdm9*^EP/EP^ spermatocytes. (F) PAS-stained sections from 3-week postpartum ovaries in wild-type and *Prdm9*^EP/EP^ mice in the B6 genetic background. Arrows show primary follicles. Bar = 100 μm. (G) Pups produced by *Prdm9*^EP/EP^ female mice. (H) Litter sizes in wild-type, *Prdm9*^-/-^ and *Prdm9*^EP/EP^ female mice. (I) Co-immunolabeling detection of RNF212 foci (red, top row), MLH1 (red, bottom row) and SYCP3 (blue) in pachytene oocyte chromatin spreads in wild-type and mutant females. White arrowheads highlight unsynapsed regions of chromosomes. Scale bars represent 10 µm. (J) Violin plot with dots showing numbers of MLH1 foci per meiotic oocyte (error bars, SEM), in wild-type, *Prdm9*^-/-^, and *Prdm9*^EP/EP^ mice. P-values were calculated using Mann-Whitney U test with Tukey’s multiple testing corrections. (K) Diagram showing proportions of PRDM9-dependent and -independent crossovers in progeny of B6WSBF2.*Prdm9*^EP/EP^ females (n = 94).

This mutation was not only hypomorphic, but also yielded an unanticipated phenotype: sexual dimorphism (Fig. S1C). Homozygous B6.*Prdm9*^*EP/EP*^ males exhibited the expected meiotic arrest and infertility, phenocopying both the *Prdm9*-null condition (8) and the methyltransferase-dead *Prdm9* allele recently reported (15) (Fig. S1C). In contrast to wild-type B6 controls, ovaries from prepubescent B6.*Prdm9*^*EP/EP*^ females were smaller but contained all developmental stages including primordial and primary follicles (Fig.1F, S1C). In contrast to the infertile males, all tested homozygous B6.*Prdm9*^*EP/EP*^ females produced at least one—and up to five—litters when mated to B6 males in fertility tests over a period of 6 months (Fig.1G-H, Tables S3 and S4). The offspring were grossly normal and healthy, and those tested—males and females—were fertile. To determine whether B6.*Prdm9*^*EP/EP*^ oocytes exhibit normal cytological features of recombination, we performed immunofluorescence staining on meiotic chromatin spreads from P0 ovaries, using antibodies against the crossover regulator RNF212, the crossover marker MLH1, and the synaptonemal complex (SC) protein SYCP3 (Fig.1I). Most (∼65%) of B6.*Prdm9*^*EP/EP*^ pachytene oocytes exhibited homologous synapsis of all chromosomes with numbers of crossovers similar to those in wild-type B6 control oocytes (Fig. 1I-J, Table S5). For comparison to *Prdm9*-deficent females, we stained P0 oocytes from B6.*Prdm9*^*-/-*^ for the crossover marker MLH1 and the SC marker SYCP3. In this case, most B6.*Prdm9*^*-/-*^ pachytene oocytes (∼67%) showed widespread asynapsis and a low number of MLH1 foci on synapsed chromosomes. However, about 4% of B6.*Prdm9*^*-/-*^ pachytene oocytes exhibited fully synapsed chromosomes stained with MLH1 foci (Fig.1I-J, Table S5). Together, these results show that female meiosis can tolerate lower levels of active PRDM9 than male meiosis, achieving sufficient recombination to ensure viable eggs. Thus, the requirement for PRDM9-dependent hotspot activation in mammalian reproduction is subject to sex-specific control. These results also provide evidence that a PRDM9-independent pathway promotes crossovers in oocytes in the absence of PRDM9.

We next investigated whether the absence of an efficient PRDM9-dependent hotspot activation mechanism promotes recombination at PRDM9-independent sites. To determine where crossover sites occur in B6.*Prdm9*^*EP/EP*^ females, we outcrossed B6.*Prdm9*^*EP*^ mice to WSB/EiJ (WSB) mice, producing B6WSBF2.*Prdm9*^*EP/EP*^ mice. As expected, B6WSBF2.*Prdm9*^*EP/EP*^ males were infertile with low testis weight (Table S6). We backcrossed three B6WSBF2.*Prdm9*^*EP/EP*^ females to wild-type B6 males, and genotyped both the F2 females and 20 of their progeny with a genome-wide SNP array (16). This allowed us to map crossovers that had occurred in the oocytes giving rise to these progeny. We excluded crossovers in the progeny that were inherited from their F2 mothers, yielding 94 informative crossovers that occurred in the *Prdm9*^*EP/EP*^ oocytes. Of these, 26 occurred in intervals with no known B6 PRDM9-dependent H3K4me3 peak (27.7%). To exclude the possibility that these 26 putative PRDM9-independent crossovers occurred at heretofore unrecognized PRDM9 binding sites on the WSB chromosomes, we searched the B6 and WSB genomic sequences in the crossover regions for PRDM9 binding motifs (17). Of these 26 intervals, 17 (18.1% of total) had no novel PRDM9 binding motifs in the WSB sequence relative to the B6 sequence, and are considered PRDM9-independent. Of the 94 total crossovers, 12 (12.8%) could be classified as clearly PRDM9-dependent based on very high-resolution crossover intervals containing a known hotspot (Fig. 1K). Interestingly, the B6WSBF2.*Prdm9*^*EP/EP*^ females produced more offspring than B6.*Prdm9*^*EP/EP*^ females (avg. offspring per female per month 4.55 vs 2.69, respectively; Table S3), suggesting an effect of genetic modifiers from the WSB genetic background, or hybrid vigor. These results provide direct evidence for PRDM9-independent recombination events occurring in oocytes that gave rise to healthy offspring.

### Genetic background and sex-limited requirement for PRDM9 in mice

The human case of a fertile PRDM9-null female (12), and the higher fertility in the B6WSBF2.*Prdm9*^*EP/EP*^ females, led us to examine the impact of different genetic backgrounds on meiotic recombination and fertility in the face of *Prdm9* deficiency. To that end, we introgressed the B6.*Prdm9*-null allele onto the CAST/EiJ and C3H/HeJ inbred backgrounds for five generations (∼5% residual heterozygosity, henceforth CAST.*Prdm9*^*-/-*^ and C3H.*Prdm9*^*-/-*^, respectively). These strains derive from distinct house mouse subspecies that diverged ∼0.5 million years ago (18, 19). Strikingly, CAST.*Prdm9*^*-/-*^ females have functional oocytes and are fertile, producing grossly healthy, fertile offspring, while C3H.*Prdm9*^*-/-*^ females, like B6.*Prdm9*^*-/-*^ females, are infertile (Fig. 2A-C, Table S4). To determine the recombination status in P0 CAST.*Prdm9*^*-/-*^ oocytes, we immunostained CAST.*Prdm9*^*-/-*^ ovarian meiotic spreads with antibodies against MLH1 and SYCP3 (Fig. 2D, Table S5). We observed normal synapsis and crossover formation in ∼ 96% of CAST.*Prdm9*^*-/-*^ pachytene oocytes, although there was a significant reduction in the average number of crossovers in these oocytes compared to the CAST.*Prdm9*^+*/*+^ control oocytes (Fig 2D-E, Table S5). Interestingly, B6CASTF1.*Prdm9*^*-/-*^ and C3HCASTF1.*Prdm9*^*-/-*^ females were fertile, while B6C3HF1.*Prdm9*^*-/-*^ females were infertile (Fig. 2C, Table S4), indicating the presence of one or more dominant modifiers in the CAST genetic background that abrogate the requirement for PRDM9 in oocytes for fertility. All *Prdm9*-deficient males of the genetic backgrounds we tested—B6.*Prdm9*^*-/-*^ and C3H.*Prdm9*^*-/-*^—exhibited meiotic arrest (Fig. S2A) and infertility (data not shown). However, we did observe round spermatids and elongating sperm in histological sections of some CAST.*Prdm9*^*-/-*^ testes (Fig. S2A-C), although these males were infertile. This suggests a rescue of meiotic arrest in some spermatocytes, as previously observed in PWD.*Prdm9*^*-/-*^ males (11). In conclusion, the requirement for PRDM9 in recombination initiation and fertility in mice is sexually dimorphic, and is modulated by background-specific genetic modifiers that lead to fertility despite PRDM9 deficiency.

**Fig. 2:**
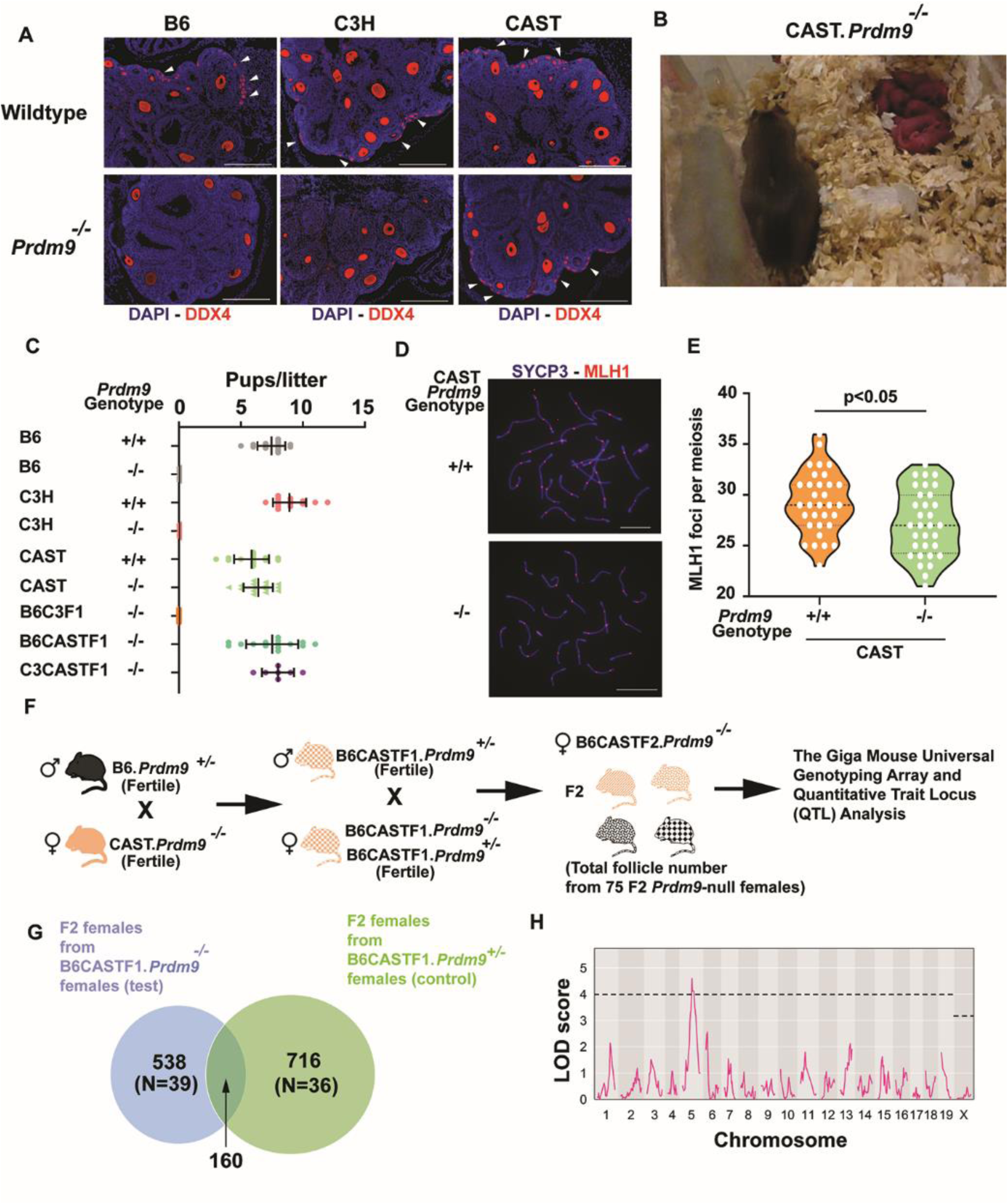
CAST/EiJ *Prdm9*-null females are fertile. (A) Immunofluorescence staining of histology sections from 3-week postpartum ovaries in mice of different genetic backgrounds. DDX4 (red, also known as MVH) marks oocytes. DNA is stained with DAPI (blue). Arrowheads indicate primordial follicles. Bar = 100 μm. Oocyte depletion in *Prdm9*^*-/-*^ mice in the B6 and C3H genetic backgrounds is seen in contrast to survival of oocytes in wild-type and *Prdm9*^*-/-*^ mice in the CAST genetic background. (B) Pups produced by CAST.*Prdm9*^*-/-*^ female mice. (C) Reproductive productivity of wild-type and *Prdm9*^*-/-*^ females in different genetic backgrounds. (D) Co-immunolabeling detection of MLH1 foci (red) in pachytene oocyte chromatin spreads, also labeled with antibody against SYCP3 (blue) in wild-type and *Prdm9*^*-/-*^ CAST females. Scale bars = 10 µm. (E) Violin plot with dots showing numbers of MLH1 foci per oocyte (error bars, SEM). The genotypes of mice tested are indicated below the graphs. P-values were calculated using Mann-Whitney U test with Tukey’s multiple testing corrections. (F) Scheme of construction of B6CASTF2.*Prdm9*^*-/-*^ female cohorts used for SNP array genotyping (GigaMUGA). Seventy-five F2 females were generated by crossing B6CASTF1.*Prdm9*^-/-^ and B6CASTF1.*Prdm9*^+/-^ females with B6CASTF1.*Prdm9*^+/-^ males. Eight-week-old B6CASTF2.*Prdm9*^-/-^ females were phenotyped for oocyte quantity and genotyped. (G) Venn diagram showing common crossover sites between F2 females generated by crossing B6CASTF1.*Prdm9*^-/-^ (test, blue) and B6CASTF1.*Prdm9*^+/-^ (control, green) females with B6CASTF1.*Prdm9*^+/-^ males. 160 common crossover sites are highlighted by overlap. Total of observed crossover sites are highlighted in the center of each circle. N represents total number of F2 females analyzed in each group. (H) Quantitative trait locus (QTL) mapping, with the total number of ovarian follicles (refer to methods for details) per B6CASTF2.*Prdm9*^-/-^ female as the phenotype. QTL (pink) reached significance on Chromosome 5, with a peak at ∼100.4 Mb (1.5 LOD drop: 72.9-127.65Mb, mm10).

In light of the PRDM9-independent crossovers we observed in B6WSBF2.*Prdm9*^*EP/EP*^ females, we questioned whether recombination hotspot usage is altered in the absence of PRDM9 in CAST females. To this end, we crossed B6CASTF1.*Prdm9*^*-/-*^ females with B6CASTF1.*Prdm9*^+*/-*^ males to generate 39 B6CASTF2.*Prdm9*^*-/-*^ females (test group) (Fig. 2F). It is expected that in these F2 test progeny more than 50% of crossovers will occur at unique (PRDM9-independent) crossover sites, given that female house mice typically have recombination rates higher than those of males. To resolve the PRDM9-dependent and -independent crossover sites, we compared the above test progeny with a group of 36 B6CASTF2.*Prdm9*^*-/-*^ F2 females generated by crossing B6CASTF1.*Prdm9*^+*/-*^ females with B6CASTF1.*Prdm9*^+*/-*^ males (control group) (Fig. 2F). These F2 control progeny should have most or all crossovers restricted to PRDM9-dependent hotspots, due to the presence of a wild-type *Prdm9* allele in both parents (Fig. 2F). We observed an overlap of only ∼30% (35% to 45% expected by random chance) between crossovers in F2 test progeny born of B6CASTF1.*Prdm9*^*-/-*^ mothers, and crossovers in F2 control progeny born of B6CASTF1.*Prdm9*^+*/-*^ mothers (Fig. 2G). The other ∼70% (378 of 538 crossovers) were unique and most likely PRDM9-independent crossovers contributed by the *Prdm9*-null mothers of the test progeny (Fig. 2G). Because crossovers are mapped at low resolution in this dataset, and because we do not have a detailed map of PRDM9-dependent hotspots in the CAST genome, it was not possible for us to classify these crossovers as *Prdm9*-dependent or -independent. Overall, these data suggest a novel set of crossover sites in B6CASTF1 oocytes in the absence of PRDM9.

To identify the CAST modifier(s) that allow *Prdm9*-null female fertility, we used the 75 B6CASTF2.*Prdm9*^*-/-*^ female mice to perform quantitative trait locus (QTL) mapping with the total number of ovarian follicles per female as the phenotype (Fig. 2F and H, see methods for details). We chose this phenotype because the number of oocytes in mature ovaries correlates with the efficacy of meiotic DSB repair and oocyte survival (20). The analysis yielded a significant QTL on Chromosome 5, with a peak at ∼100.4 Mb (1.5 LOD drop: 72.9-127.65Mb) (Fig. 2H, S2D-E). Intriguingly, this QTL contains two critical meiotic genes: ring finger protein 212 (*Rnf212* at 108.7Mb) and checkpoint kinase 2 (*Chk2* at 110.8Mb). The success of the meiotic recombination process depends on the efficient repair of meiotic DSBs and crossover formation, while oocyte survival depends on successful passage through a checkpoint that monitors DNA damage. The roles of RNF212 and CHK2 in DNA damage surveillance in oocytes make them possible candidates that might act as genetic modifiers of PRMD9 (20, 21). RNF212 is a SUMO ligase essential for crossover formation, and also mediates oocyte quality control (21). It has been reported that localization of RNF212 to DSB sites acts as a “memory” of unrepaired DSBs, thus promoting apoptosis of defective oocytes during the diplotene to dictyate meiotic substage transition (21). Although loss of *Rnf212* is compatible with survival of oocytes in spite of persistent DSBs and synapsis defects, fertility is not rescued in female *Rnf212* knockout mice (21). The other candidate gene, *Chk2*, encodes a meiotic checkpoint kinase responsible for DNA damage surveillance in oocytes (20). Ablation of CHK2 prevents oocyte elimination in response to both radiation-induced DNA damage and persistent meiotic DSBs due to genetic mutation in *Trip13* (20). Both the *Rnf212* and *Chk2* genomic sequences are well conserved between B6 and CAST mice (Fig. S3A-B); however, regulatory variants outside the genes themselves may play a critical role.

### Modulation of a meiotic DNA damage checkpoint permits female-limited fertility in Prdm9-null B6 mice

The first candidate modifier gene we considered was *Rnf212*. In meiotic spreads, RNF212 protein expression and co-localization patterns were similar between CAST.*Prdm9*^*-/-*^ meiotic oocytes and CAST.*Prdm9*^+/+^ oocytes (Fig. S4). As mentioned above, crossover formation in CAST.*Prdm9*^*-/-*^ meiotic oocytes is normal (Fig. 2D-E), suggesting that the role of RNF212 in crossover formation in CAST.*Prdm9*^*-/-*^ remains intact. While the oocyte count of B6.*Rnf212*^−/−^ females is normal, these mice are infertile due to an absence of crossovers (21). RNF212 deficiency promotes survival of oocytes with genetic and radiation-induced DNA damage, suggesting an additional role for RNF212 as a pro-apoptotic cell-cycle regulator that promotes elimination of defective oocytes (21). In CAST.*Prdm9*^*-/-*^ females, delayed activation of RNF212 in this capacity could conceivably promote the survival of oocytes with unrepaired DSBs, while still fulfilling its role in crossover formation. Further investigation is needed to understand the exact role of RNF212 in PRDM9-dependent and -independent meiotic recombination in CAST oocytes. However, because of the similarities between *Prdm9*^+/+^ and *Prdm9*^*-/-*^ CAST oocytes in both in RNF212 expression and RNF212 co-localization, we consider it an unlikely modifier-gene candidate.

To determine directly whether *Chk2* deficiency allows *Prdm9*-null oocytes to complete meiosis, we generated *Prdm9*^*-/-*^*Chk2*^*-/-*^ double knockout females in the B6 genetic background (henceforth, B6.*Prdm9*^*-/-*^*Chk2*^*-/-*^). Among P0 oocytes from B6.*Prdm9*^*-/-*^ single mutants, most exhibited widespread asynapsis and persistent unrepaired DSBs, revealed by pervasive BRCA1, γH2AFX, and IHO1 signals (Fig. S5A-C). Only 4% of these P0 oocytes had fully synapsed chromosomes stained with RNF212, MLH1, and MLH3 foci (markers of mature crossovers) (Fig. 3A-D, S5D, Table S5). This phenotype was substantially improved in B6.*Prdm9*^*-/-*^*Chk2*^*-/-*^ double-mutant oocytes, with a significant increase in the number of oocytes with fully synapsed chromosomes devoid of BRCA1, γH2AFX, and IHO1 (∼26%, Fig. S5A-C, E). We also observed a ∼threefold increase in the number of dictyate oocytes in P0 B6.*Prdm9*^*-/-*^*Chk2*^*-/-*^ double-mutant females, compared to B6.*Prdm9*^*-/-*^ females (Fig. S5A-D, F). Only a small number of growing follicles survived in B6.*Prdm9*^*-/-*^ females (Fig. 3G-H). We were able to isolate metaphase I oocytes from young B6.*Prdm9*^*-/-*^ and B6.*Prdm9*^*-/-*^*Chk2*^*-/-*^ females; here we noted a significant increase (∼25% increase) in the number of metaphase oocytes with normal chiasmata between the homologous chromosomes in B6.*Prdm9*^*-/-*^*Chk2*^*-/-*^ females relative to B6.*Prdm9*^*-/-*^ females (Fig. 3E-F). We conclude that CHK2 eliminated oocytes that failed to repair meiotic DSBs, but those that completed recombination were not eliminated and were able to progress through folliculogenesis. Although 4% of B6.*Prdm9*^*-/-*^ oocytes at P0 had a full complement of crossovers, this is probably not enough for fertility, especially given further oocyte elimination after P0 during follicle formation and atresia. We propose that the removal of *Chk2* allowed more *Prdm9*^*-/-*^ oocytes to escape the meiotic DNA damage checkpoint and subsequently complete DSB repair, resulting in a higher oocyte reserve.

**Fig. 3:**
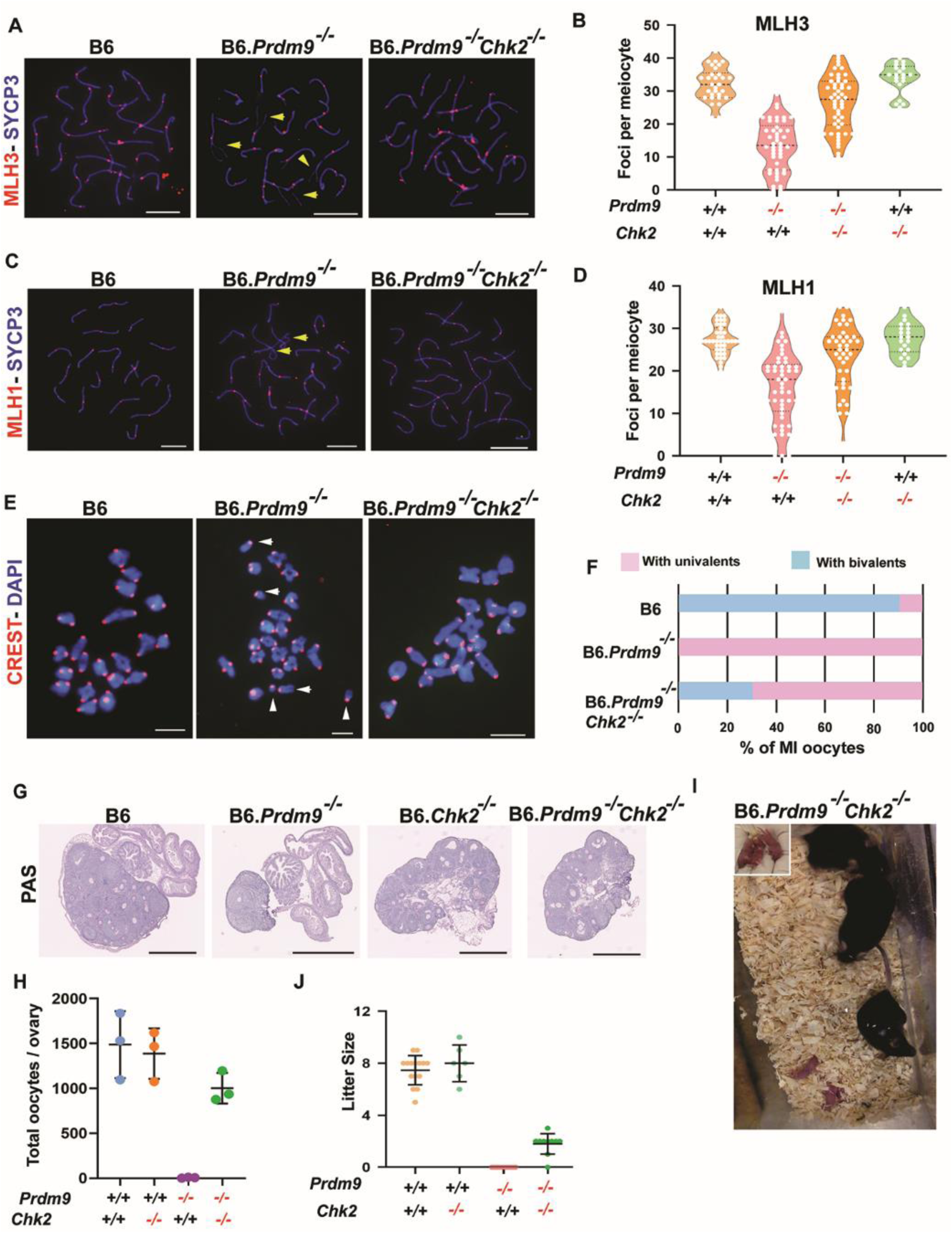
*Chk2* ablation rescues fertility in *Prdm9*-null females in the B6 genetic background. (A-D) Co-immunolabeling detection of MLH3 foci (red, A), MLH1 (red, C) and SYCP3 (blue, both A and C) in pachytene oocyte chromatin spreads from wild-type and mutant females. Yellow arrowheads highlight unsynapsed regions of chromosomes. Scale bars = 10 µm. Violin plot with dots showing numbers of MLH3 (B) and MLH1 (D) foci per meiotic oocyte (error bars, SEM). The genotypes of mice tested are indicated below the graphs. p values were calculated using Mann-Whitney U test with Tukey’s multiple testing corrections. (E-F) Chromosome configuration in meiosis I metaphase analyzed by chromosome spreads from different mutant and control oocytes (E) and quantification (F). In E, DNA (blue) and kinetochores (red) were detected by DAPI and CREST antiserum, respectively. Scale bars = 10 μm. Multiple univalents in a B6.*Prdm9*^*-/-*^ oocyte are indicated with white arrowheads. The examples of oocytes from B6 and B6.*Prdm9*^*-/-*^ *Chk2*^*-/-*^ mice reveal all chromosomes organized in bivalents. (G) PAS-stained histological sections of 3-week postpartum ovaries from mice of different genetic backgrounds. Bar = 200 μm. (H) Oocyte quantification in mutant and control animals. (I) Pups produced by B6.*Prdm9*^*-/-*^ *Chk2*^*-/-*^ female mice. (J) Litter sizes in mutant and control females.

To test the impact of improved meiotic phenotypes on oocyte number and fertility, we estimated the oocyte number in B6.*Prdm9*^*-/-*^*Chk2*^*-/-*^ females and performed fertility tests with wild-type B6 males. In contrast to B6.*Prdm9*^*-/-*^ females, B6.*Prdm9*^*-/-*^*Chk2*^*-/-*^ females produced at least one litter of grossly healthy, fertile offspring over a six-month period (Fig. 3I-J, S6A, Table S4), indicating some functional ovarian reserve. However, there was a high incidence of pregnancy losses and perinatal deaths (Fig. S6B), suggesting that oocyte quality is compromised, consistent with the observations of metaphase oocytes in B6.*Prdm9*^*-/-*^*Chk2*^*-/-*^ females. The extent of this fertility was much lower than that observed in CAST.*Prdm9*^*-/-*^ females. In contrast to females, *Chk2* deficiency did not rescue fertility in B6.*Prdm9*^*-/-*^ males; both single- and double-mutant males were infertile with complete meiotic arrest (Fig. S5G). Together, these results demonstrate that, in females, a *Prdm9*-independent recombination pathway can give rise to the required number of crossovers for production of healthy offspring, facilitated by inactivation of the *Chk2*-dependent DNA damage surveillance mechanism.

## Discussion

In this study, we find that female meiosis tolerates critically low levels of catalytically active PRDM9, or even no PRDM9 at all in some genetic backgrounds, because a PRDM9-independent recombination pathway can compensate successfully. The meiotic DNA-damage repair protein CHK2 acts as a modifier, with its modification or abrogation allowing completion of meiosis and fertility in females lacking active PRDM9. This is not the case for males; *Prdm9*^*EP/EP*^ and *Prdm9*-null males in all genetic backgrounds examined were infertile. Overall, these results demonstrate that the requirement for functional PRDM9 during meiosis varies substantially by genetic background as well as by sex, and that there are alternative pathways for effective recombination.

As shown by our genetic analyses, modification or even abrogation of the CHK2-dependent DNA damage checkpoint activation is a critical determinant of female fertility in the absence of PRDM9. CHK2 triggers elimination of oocytes with persistent DNA damage. It is possible that PRDM9-independent DSB repair is inefficient or prolonged, which would cause DSBs to persist past the timely activation of the CHK2-dependent DNA damage checkpoint, leading to oocyte elimination. Ablation of *Chk2* and consequent elimination of checkpoint surveillance may promote survival of oocytes by allowing extra time for PRDM9-independent crossovers to accumulate, permitting a sufficient number of oocytes to complete meiosis. However, this is not a fully adequate explanation of the modifier effect, because although B6.*Prdm9*^*-/-*^*Chk2*^*-/-*^ females were fertile, they were not as productive as CAST.*Prdm9*^*-/-*^ females. Thus, while CHK2 is clearly mechanistically involved, it does not completely explain the disparity between B6 and CAST female fertility in the absence of PRDM9. Furthermore, the predicted causative CAST variant(s) in *Chk2* seem to be dominant and possibly neomorphic, unlike the recessive null mutation that restores fertility in B6.*Prdm9*^*-/-*^*Chk2*^*-/-*^ females. It is therefore possible that both *Chk2* and *Rnf212* are modifiers of the PRDM9-independent fertility phenotype, or that additional CAST genome variants affect timing or efficacy of DNA repair, the checkpoint, or other features of meiosis that determine whether or not oocytes activate the checkpoint. It may also be the case that repair of PRDM9-independent DSBs is more efficient in the CAST genetic background than in the B6, with fewer oocytes exhibiting a number of DSBs sufficient to activate DNA damage checkpoint. Ability to repair DSBs may be a limiting factor in B6 mice; the high pre- and perinatal death rate (Fig. S6B) and the aneuploidy rate we observed in developing oocytes (Fig. 3E and F) indicate that, in B6 mice, the PRDM9-independent pathway leads to maturation of defective oocytes. Although similar modification of an infertility phenotype via removal of *Chk2* was reported for *Trip13-*deficient females (20), there are critical differences between causes of infertility in *Trip13-* and *Prdm9-*deficient females. In *Trip13* mutants, crossovers originate from PRDM9-dependent DSBs, and complete synapsis is achieved (20, 22); in contrast, *Prdm9*-deficient oocytes exhibit widespread chromosomal asynapsis, and form crossovers at ectopic DSBs. *Trip13*-deficient oocytes are eliminated due to inefficient repair of DSBs in a non-crossover pathway (20). Removal of *Chk2* in *Trip13* mutants reestablishes ovarian reserve and fertility by allowing oocyte survival and additional time for DSB repair (20); however in *Prdm9*-deficient oocytes, *Chk2* removal must also allow chromosomal synapsis and recombination at ectopic, non-PRDM9 DSBs. To our knowledge, this is the first evidence that modulation of a DNA damage checkpoint protein can allow survival of oocytes that undergo genetic recombination via a PRDM9-independent pathway.

While checkpoint modulation appears to be a feature allowing oocyte, but not spermatocyte, survival in the absence of PRDM9, at least in certain genetic backgrounds, the question of the genetic and cellular mechanisms behind this dimorphism is undoubtedly more complex. Although recombination is essentially the same process in both sexes, spermatogenesis and oogenesis are fundamentally different cell differentiation programs. Two key differences unique to males that might explain the sexually dimorphic requirement for PRDM9 are meiotic sex chromosome inactivation (MSCI), which is the silencing of most genes on the sex chromosomes from pachynema of meiotic prophase I into spermiogenesis (23), and deployment of Piwi-interacting RNAs (piRNAs) (24, 25). MSCI is triggered by XY asynapsis in spermatocytes and is marked by sequestration of the sex chromosomes in the ‘sex body’ (23). Spermatocytes undergoing PRDM9-deficient meiotic arrest in B6 mice do not form a normal sex body, as unsynapsed autosomes and unrepaired DSBs compromise recruitment of silencing factors to sex chromosomes (8, 26, 27). This could potentially lead to failure of MSCI, triggering meiotic arrest of PRDM9-deficient spermatocytes (8, 26). Inappropriate critical gene silencing on the asynapsed autosomes due to meiotic silencing of unsynapsed chromatin (MSUC) may also contribute to spermatocyte elimination (23). Oocytes, which are affected by MSUC to a lesser extent, may escape these outcomes (23). In mammals, piRNAs are necessary to suppress expression of L1 retrotransposons during spermatogenesis (24, 25). Mutation or deficiency of the piRNA pathway contributes to male-limited sterility (28-30), and it was recently reported that the expression of piRNAs is misregulated in *Prdm9*-null spermatocytes (31). Thus, it is possible that dysregulation of the piRNA pathway contributes to the post-meiotic problems in *Prdm9*-null germ cells that undergo partial meiotic arrest, as seen in PWD (11), CAST, and other genetic backgrounds (this report).

In addition to invoking intriguing meiotic mechanisms, the sexually dimorphic responses to PRDM9 deficiency described here have significant evolutionary implications. *Prdm9* is the first and only known mammalian speciation gene (11). *Prdm9* interacts with an unknown element on the X chromosome to cause male-limited meiotic arrest and hybrid sterility in F1 hybrids between the females of the *M. m. musculus* strain PWD/PhJ, and the males of certain *M. m. domesticus* strains, including B6 (32, 33). Sterile (PWDXB6)F1 hybrid males exhibit a significant enrichment of DSBs at PRDM9-independent hotspots and high rates of autosomal asynapsis that trigger pachytene checkpoint activation (33, 34). Chromosome asynapsis is also observed in F1 hybrid females, although the phenotype is markedly less severe than in F1 hybrid males, and females remain fertile (33, 35). Our findings suggest that F1 hybrid females may retain fertility by effectively utilizing a PRDM9-independent DSB repair mechanism to evade checkpoint activation. Our work further nominates *Chk2* as a key modifier of these sex and strain differences in the meiotic tolerance for PRDM9-independent DSB repair.

The pattern of male-limited hybrid sterility in the (PWDxB6)F1 model follows Haldane’s rule, which postulates that when only one sex of an interspecies hybrid experiences sterility or inviability, it is the heterogametic sex (36, 37). Many hypotheses, largely based on role of sex chromosomes, have attempted to explain Haldane’s rule, including the faster-male theory, the faster X theory, meiotic drive, and the dominance theory (38). However, the mechanistic basis for sexually dimorphic hybrid sterility remains unclear in most documented cases. The sexually dimorphic requirement for PRDM9 for fertility and the relaxed stringency of DNA damage surveillance in females that we report here provide a possible explanation for Haldane’s rule in the (PWDXB6)F1 hybrid sterility model. Notably, although hybrid sterility in this system depends on a genetic interaction between *Prdm9* and an X-linked locus (33, 35), the sex-specific manifestation of this sterility appears to be driven, at least in part, by sex and genetic differences at an autosomal gene, *Chk2*.

Finally, *Prdm9* is a fundamental evolutionary innovation that is thought to have evolved to direct recombination away from functional elements, safeguarding these regions from recombination-associated mutagenesis (7). As in all cases of evolutionary innovation, however, there are trade-offs. Use of a DNA-binding protein to specify regions of recombination requires that sufficient numbers of the cognate binding sites of that protein remain intact. However, PRDM9-dependent hotspots can extinguish themselves via gene conversion, leading to gradual erosion of the hotspot-binding motifs recognized by a particular PRDM9 variant and, ultimately, meiotic failure and infertility (39, 40). One mechanism through which populations may avoid this fate is the emergence of new *Prdm9* alleles that recognize new suites of hotspots (39, 40). Indeed, *Prdm9* is highly polymorphic and its DNA-binding domain is rapidly evolving via positive selection (1). Our findings introduce an additional, novel complexity to the dynamic interplay between hotspot erosion and reproductive fitness; i.e., sex differences in the usage or effectiveness of PRDM9-independent pathways may render one sex more resilient to the loss of PRDM9 activity, leading to potential sex-specific fitness effects of hotspot erosion. Further work is required to elucidate the mechanistic details of the sexually dimorphic responses to lack of PRDM9; nevertheless, the theoretical implications raised by this phenomenon present fascinating new insights into the checks and balances that constrain one of the most fundamental evolutionary processes in mammals—genetic recombination.

## Acknowledgments

We thank all members of the Bolcun-Filas, Handel, Paigen and Petkov labs for insightful discussions and suggestions. We acknowledge the expertise and contributions of the Jackson Laboratory Scientific Services, including the Histology core, the Microscopy core, Genome Technologies, Genetic Engineering Technology and Mouse Resources for their expertise and help during this project. We thank the Knockout mouse project (KOMP2) at the Jackson Laboratory for providing mice. We thank Drs. Attila Toth, Paula Cohen and Neil Hunter for sharing their antibody resources;

## Author contributions

Conceptualization, T.B. and N.R.P; Methodology, T.B., N.R.P., B.L.D and E.B-F.; Investigation, T.B., N.R.P.,B.L.D; C.E, R.A.L., and C.B.; Writing – Original Draft, T.B and N.R.P.; Writing – Review & Editing, T.B., N.R.P.,B.L.D; C.E, R.A.L., C.B; K.P; E.B-F; P.M.P and. M.A.H; Funding Acquisition, T.B., N.R.P.,B.L.D;. K.P; E.B-F; P.M.P and. M.A.H; Resources, T.B, N.R.P, B.L.D; K.P; E.B-F; P.M.P and M.A.H.; Supervision, T.B, B.L.D; E.B-F; P.M.P and M.A.H.

## Data and materials availability

All data is available in the main text or the supplementary materials. All data, code, and materials used in the analysis are available in some form to any researcher or commercially for purposes of reproducing or extending the analysis. H3K4me3 ChIP seq data are available at NCBI Gene Expression Omnibus (GEO; http://www.ncbi.nlm.nih.gov/geo) under accession number GSE144144.

