## Supplemental data for "Sexual dimorphism in the meiotic requirement for PRDM9: a mammalian evolutionary safeguard"

**This PDF file includes:**

Materials and Methods
Figs. S1 to S6
Tables S1 to S6

### Materials and Methods

#### *Ethics Statement*

The animal care rules used by The Jackson Laboratory are compatible with the regulations and standards of the U.S. Department of Agriculture and the National Institutes of Health. The protocols used in this study were approved by the Animal Care and Use Committee of The Jackson Laboratory (Summary #04008 and 15001). Euthanasia was carried out by cervical dislocation.

#### *Mouse Strains*

Mice used in this study were acquired from The Jackson Laboratory. The following strains were used: C57BL/6J (stock number 000664), WSB/EiJ (stock number 001145), CAST/EiJ (stock number 000928), C3H/HeJ (stock number 000659), B6;129P2-*Prdm9*$^{tm1Ymat}$/J (stock number 010719), C57BL/6J-Prdm9$^{<em2Kpgn>}$/Kpgn (stock number 28854; *Prdm9*$^{EP}$). C57BL/6N-A$^{tm1Brd}$ *Chek2*$^{tm1b(EUCOMM)Hmgu/}$JMmucd (stock number 047090-UCD, also known as *Chk2*) mice were acquired from the KOMP repository at the Jackson Laboratory. The B6;129P2-*Prdm9*$^{tm1Ymat}$/J mice were backcrossed to C57BL/6J for 10 generations before use in this study. C3H-*Prdm9*$^{-/-}$ and CAST/EiJ-*Prdm9*$^{-/-}$ congenic mice were generated by backcrossing with B6;129P2-*Prdm9*$^{tm1Ymat}$/J mice for five generations. C57BL/6J-Prdm9$^{<em2Kpgn>}$/Kpgn were generated by introducing point mutations (glu365pro) via CRISPR/Cas9 gene editing, onto the C57BL/6J genetic background (Fig. S1A). Gene editing was performed by the Genetic Engineering Technology core at the Jackson Laboratory.

#### *Fertility tests*

Control and experimental females (2-6 months old) were housed with fertile males for a period of up to 6 months. A female was considered fertile if she gave live birth at least once. Litter size was determined by counting pups on the day of birth. Embryo loss during pregnancy was assessed by comparing the number of live births per pregnancy. Briefly, females were mated to wild-type males and the presence of a copulation plug was checked the next morning and recorded. At 19.5 dpc (days post coitum), pregnant females were expected to deliver. Embryo loss during pregnancy was clear due to sinking of stomach in pregnant females due to reabsorption of embryos by 16.5 dpc.

#### *Tissue Fixation, Histology, immunofluorescence and follicle quantification*

Whole testes and ovaries were fixed for 24h in Bouin's fixative at room temperature, then washed three times with 70% ethanol at room temperature for 1h per wash. Testis cross-sections were stained with Periodic acid-Schiff-diastase (PAS) by standard methods, and imaged at 40X magnification with a Nanozoomer 2.0HT. Ovaries from 3- and 8-week-old females were fixed and embedded in paraffin, serially sectioned at 5μm and stained by hematoxylin and eosin. Follicle quantification was performed as published previously (*1*). In brief, every fifth section was examined for the presence of the following classes of oocytes/follicles: primordial, primary, secondary, and mature. For quantitative trait locus (QTL) analysis, total follicle number was used. Statistical differences in follicle number between different genotypes were evaluated using the non-parametric two-tailed Mann-Whitney test using GraphPad Prism 7.

For immunofluorescence, testes from 2-month-old and ovaries from 3-week-old mice were fixed in 4% PFA and sectioned at 5 μm thickness, matured overnight, de-waxed, rehydrated, and heated in 10 mM sodium citrate buffer (pH 6.0) for antigen retrieval. Slides with testis sections were incubated with ADB buffer (1.5% BSA, 5% donkey serum, 0.05% Triton X-100 in PBS, 0.2x cocktail of protease inhibitors) for 1 h at room temperature. Slides were incubated overnight with diluted anti-PRDM9 (1:100) (*2*), anti-γH2AFX (1:5000, Upstate #07-164) and anti-SYCP3 (1:100, D-1; Santa Cruz #74569) in ADB buffer. Slides with ovary sections were stained using rabbit anti-MVH (1:500, Abcam #27591). Immunofluorescent staining was performed by staining the sections for 2 h at 37°C, using diluted appropriate secondary antibodies conjugated with Alexa 488/FITC, Alexa 594/CY3 and Alexa 647/CY5 (Molecular Probes/Invitrogen or Jackson Immunoresearch Laboratories 1:300). Stained sections were mounted using ProLong Gold Antifade Mountant with DAPI (P36935; Molecular Probes/Invitrogen) at 4°C overnight and imaged after 24 h using a Leica SP5 confocal microscope and/or Zeiss AxioImager.Z2 epifluorescence microscope. Images were processed and adjusted using Adobe Photoshop (Adobe Systems).

***Immunostaining of spread meiocytes***

The meiocyte spreads were prepared by using the hypotonic protocol as described previously (*3*). The nuclei were immunostained using the following primary antibodies: rat polyclonal anti-SYCP3(*4*), mouse monoclonal anti-MLH1 (abcam # 14206), mouse monoclonal anti-γH2AFX (Upstate, #05-636), mouse monoclonal anti-SYCP3(D-1) (Santa Cruz #74569), guinea pig polyclonal anti-IHO1 (*5*), rabbit polyclonal anti-RNF212 (*6*), rabbit antibody anti-MLH3 (*7*) and rabbit polyclonal anti-BRCA1(C-20) (Santa Cruz # 642); and the following secondary antibodies: goat anti-Rabbit IgG-AlexaFluor488 (Molecular Probes, A -11034), goat anti-Mouse IgG-Alexa Fluor 568 (Molecular Probes, A-11031), goat anti-Rabbit IgG-Alexa Fluor 568 (Molecular Probes, A-11036), goat anti-Mouse IgG-Alexa Fluor 350 (Molecular Probes, A- 21049), goat anti-Mouse IgG-Alexa Fluor 647 (Molecular Probes, A-21236), goat anti-Rabbit IgG-Alexa Fluor 647 (Molecular Probes, A-21245) and goat anti-Guinea pig IgG-Cy3 (Chemicon, #AP108C). Stained meiocytes were mounted using ProLong Gold Antifade Mountant with DAPI (P36935; Molecular Probes/Invitrogen) at 4°C overnight and imaged after 24 h using a Zeiss AxioImager.Z2 epifluorescence microscope. Images were processed and adjusted using Adobe Photoshop (Adobe Systems). MLH1 foci were counted using Image J software (http://rsbweb.nih.gov/ij/). Statistical differences in MLH1 and MLH3 focus number per meiocyte between different genotypes were evaluated using the non-parametric two-tailed Mann-Whitney test using GraphPad Prism 7. Meiocytes with partially asynapsed or desynapsed chromosomes were also considered as a data point for the MLH1 and MLH3 analysis.

***GigaMUGA Genotyping and QTL analysis***

GigaMUGA genotyping was carried out by Neogen's commercial service. Invariant SNPs, SNPs with erroneous or missing calls in parental and F1 control samples, SNPs with >10% missing data across all samples, and sites that deviated from Hardy-Weinberg expectations were excluded. Putative genotyping errors were identified as tight double recombinants and recoded as missing data. A total of 54,629 genotypes survived these filters. Cleaned genotypes were then down-sampled to every $10^{th}$ marker to eliminate a large number of uninformative markers, reduce the impact of genotyping error on map inflation, and expedite the process of map construction. This thinned genotype dataset was then used to construct an empirical genetic linkage map with the *est_map* call in R/qtl2 (*8*). Recombination fractions were converted to map distances using the Carter-Falconer mapping function (*9*) and assuming a 1% residual genotyping error rate.

Single QTL mapping of recombinant B6CASTF2.*Prdm9*$^{-/-}$ females was performed using the linear mixed model framework implemented in the R/qtl2 package. Conditional genotype probabilities were calculated from the downsized marker dataset without the inclusion of any pseudomarkers. The non-random genetic structure among samples was specified via a kinship matrix computed using the leave-one-chromosome-out method. QTL significance thresholds were determined by 1000 permutations of the data. To improve fit to normality, follicle counts were natural log transformed prior to mapping.

***Metaphase I Spreads***

MI spreads and staining were done as previously described (*10*). Briefly, pregnant mare's serum gonadotropin (PMSG) (5 IU) was injected to 4-week-old female mice 48 hours before time of collection of oocytes. Cumulus-oocyte complex or single GV oocytes were collected in MEM/PVP medium (Minimal essential medium (MEM), 25 mM Hepes, 3 mg/ml Polyvinylypyrollidone, 2.5 uM milrinone). After removing cumulus cells, denuded oocytes were matured for Metaphase I (MI) by incubating in a single drop of CZB/glutamine (81.62 mM NaCl, 4.83 mM KCl, 1.18 mM $KH_2PO_4$, 1.18 mM $MgSO_4$ $7H_2O$, 25.12 mM $NaHCO_3$, 1.7 mM CaCl $2H_2O$, 31.3 mM sodium lactate, 0.27 mM sodium pyruvate, 0.11 mM EDTA, 3 mg/mL BSA, 7 mM Taurine, 10 ug/ml Gentamicin, 10 ug/ml Phenol red, 1 mM Glutamine). Zona pellucida was removed from matured MI oocytes by using EmbryoMax Acidic Tyrodes solution (Millipore) and then oocytes were placed in spread solution ($H_2O$, 0.16% Triton-X100, 6 mM DTT, 0.64% PFA). After slides were completely dried, slides were Incubated with blocking solution (3% BSA in PBS) for 10 min, at RT. Immunostaining was performed by incubating slides in anti-CREST antibody (Antibodies Inc (15-234-0001)) diluted in blocking buffer for 3 hrs.

After washing with blocking buffer, secondary antibody (anti-human Alexa 647) incubation was performed for 1.5 hours. Slides were mounted in Vectashield with DAPI.

#### *H3K4me3 ChIP-seq*

ChIP-seq for H3K4me3 was performed with spermatocytes isolated from 14dpp *Prdm9*$^{EP/EP}$ animals, as previously reported (*11*), using a commercially available polyclonal α-H3K4me3 antibody (EMD Millipore cat#07-473). DNA samples were sequenced on an Illumina NextSeq, with 75bp reads, and trimmed for quality using trimmomatic. Sequence data were aligned to the mouse mm10 genome using BWA v0.7.9a, and duplicate reads and reads which failed to align to unique positions in the genome were discarded. This resulted in a total of 32,894,681 aligned reads. Sequence data are available at NCBI Gene Expression Omnibus (GEO; http://www.ncbi.nlm.nih.gov/geo) under accession number GSE144144. PRDM9-dependent H3K4me3 peak locations for B6 were previously described by Baker et al. (*11*); files with these hotspot locations are available under GEO accession number GSE52628. For the present analysis, their positions were converted to mm10 using the UCSC Genome Browser tool LiftOver; this file is available under GEO accession number GSE144144.

For the wild-type B6 sample, the sequence data used were previously reported by Baker et al. (*11*). Data are available under GEO accession number GSE52628 (sample accession numbers GSM1273023 and GSM1273024). These data were mapped to mm10 using the above procedure, resulting in 44,329,881 total aligned reads.

#### *H3K4me3 Peak Calling*

H3K4me3 peaks were called using MACS v1.4, default parameters, with a p-value cutoff of 0.01, using treatment and control samples. For a control sample, the two input B6 biological replicates from Baker et al. (*11*) were merged and aligned to mm10 via the above procedure, for a total of 76,163,775 aligned reads (GEO accession number GSE52628, sample accession numbers GSM1653213 and GSM1653215). To determine overlap between known PRDM9-dependent H3K4me3 peaks and ChIP-seq peak sets, we used Bedtools (v2.27.0) intersect with default parameters, except for a requirement for 20% overlap (f = 0.20).

#### *MA Plots*

To generate MA plots, we first counted the number of sequencing reads within each peak using Bedtools (v2.27.0) coverage, employing the -counts function with default parameters. We then normalized these counts to reads per million (RPM) mapped reads, and used the normalized counts as input for MA plots.

The actual plots were generated using the plotMA function of the R package limma (https://www.r-project.org/), with default parameters.

#### *B6xWSB-Prdm9$^{EP/EP}$ Crossover Analysis*

Three (B6xWSB)F2-*Prdm9*$^{EP/EP}$ females and 20 of their ((B6xWSB)F2xB6)F3 offspring were genotyped on the GigaMUGA SNP array. Crossovers were called based on informative SNPs between B6 and WSB, using an in-house R script (available upon request). Minimum size for a genetic interval was five consecutive informative markers. Individual chromosomes with more than five crossovers were excluded from analysis. Crossovers present in the mothers were subtracted from those in the offspring. The 94 informative offspring-specific crossover intervals were assessed for overlap with known hotspots—defined as PRDM9-dependent H3K4me3 peaks in the B6 genetic background (n=18,838) (*11*)—using bedtools intersect (v2.27.0) with default parameters.

The WSB sequences (ENSEMBL) of crossover intervals with no B6 hotspot were searched for 11bp PRDM9$^{Dom2}$ binding motifs using the MEME suite tool FIMO. The PRDM9$^{Dom2}$ motif table was based on the empirically determined consensus motif (*12*). The WSB and B6 motif lists were compared, and those WSB intervals with no novel PRDM9$^{Dom2}$ binding motifs relative to the cognate B6 intervals were noted as such.

#### *Testis extract preparation and western blotting*

Crude testis extract was prepared as reported previously (*13*) from 12dpp male mice with the denoted genotypes. Samples were subjected to standard SDS-PAGE and western blotting for detection of PRDM9

(1:1000) (*2*). Subsequently, the blot was stripped and re-probed with mouse anti-β-tubulin primary antibody (1:10,000, Sigma Cat# T4026). Both primary antibodies were detected with HRP-conjugated anti-mouse secondary antibody (1:20,000, Bio-Rad Cat# 170-6516).

***Sequence alignment analysis***
The DNA sequence for *Rnf212 and Chk2* for mm10 reference, CAST/EiJ, and C57BL/6NJ were obtained from published sequence datasets at https://www.sanger.ac.uk/science/data/mouse-genomes-project and http://useast.ensembl.org/Mus_musculus/Info/Index . The alignment was performed using Blastn. Data were visualized using dot matrix plots.

***Quantification and Statistical Analysis***
Statistical tests were performed using GraphPad Prism version 7.0 and R statistical packages.

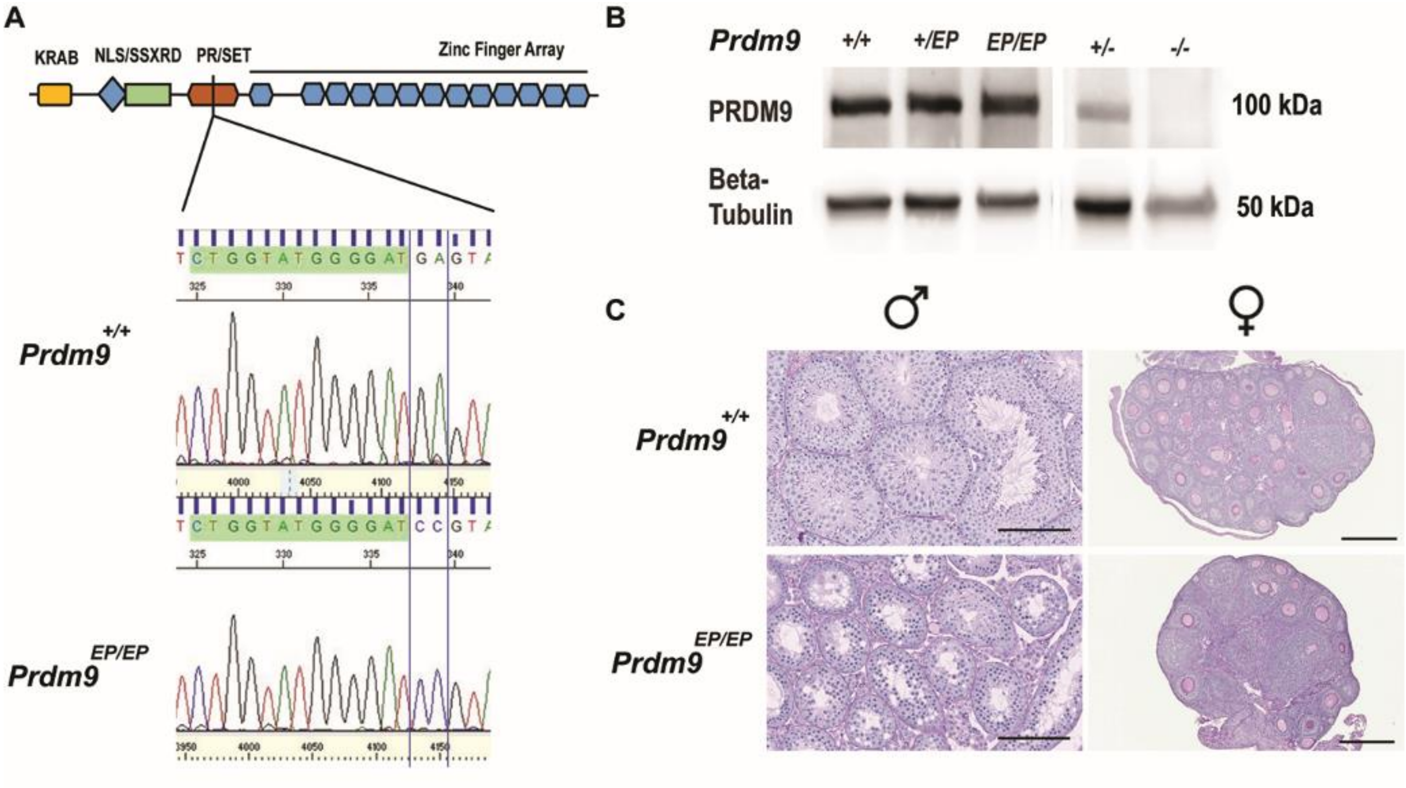


**Fig. S1.** Characterization of the *Prdm9*$^{EP}$ mutants. (A) Electropherograms showing the context of the *Prdm9*$^{EP}$ mutation within the PRDM9 PR/SET domain. (B) Western blot showing PRDM9 expression in 12 days post-partum testes isolated from wild-type, *Prdm9*$^{+/EP}$, *Prdm9*$^{EP/EP}$, *Prdm9*$^{+/-}$and *Prdm9*$^{-/-}$mice (B6 genetic background). The blot was stripped and re-probed for beta-tubulin as a loading control. (C) PAS-stained histological sections showing adult testes and 3 weeks post-partum ovaries isolated from wild-type and *Prdm9*$^{EP/EP}$ mice (B6 genetic background).

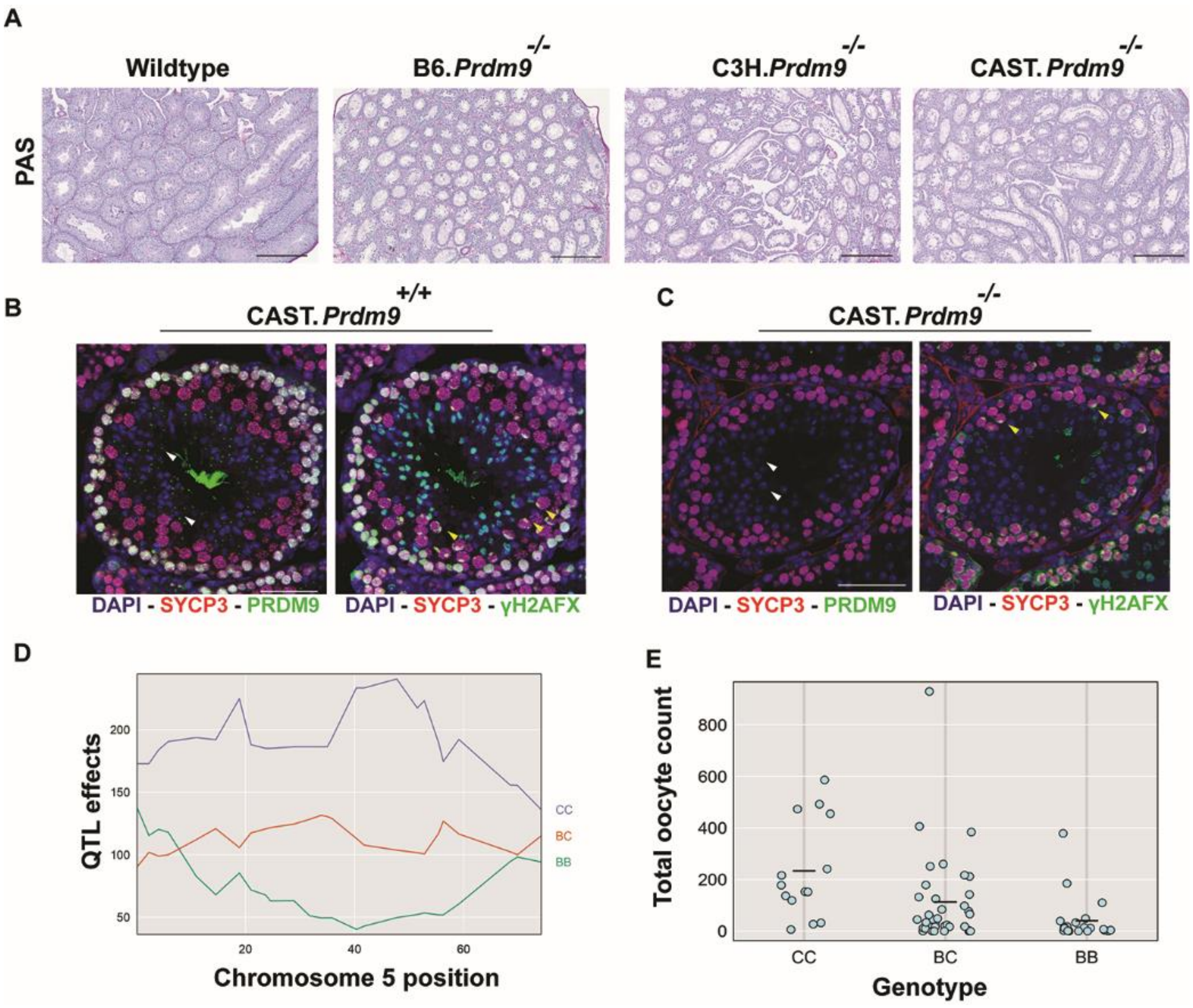


**Fig. S2.** Sexually dimorphic phenotypes of *Prdm9* deficiency in the CAST/EiJ genetic background. (A) PAS-stained histological sections of mouse testis sections of the indicated genotypes at 8 weeks of age, revealing poor progress of spermatogenesis in absence of PRDM9. Scale bars =100 µm. (B - C) Immunofluorescence staining of histological sections of CAST wild-type (B) and *Prdm9*$^{-/-}$ (C) testes stained with anti-SYCP3 (red), anti-PRDM9 (green, left panel) and anti- γH2AFX (green, right panel) antibodies; DNA is stained with DAPI (blue). The wild-type testes co-express SYCP3, and PRDM9 in pre-leptotene and leptotene cells, and in zygotene (-like) spermatocytes (B). *Prdm9*$^{-/-}$ mutants lack PRDM9 expression confirming null mutation (C). Presence of round spermatids (white arrowheads) and normal XY body (also called sex body, green) stained by γH2AFX (yellow arrowheads) suggest PRDM9 is not essential for meiosis in CAST testes (C). Scale bars =100 µm. (D & E) QTL effects and phenotype distribution plots. QTL is due to the positive effect of CAST alleles on oocyte number as highlighted by CC (violet) and BC (orange) lines, while B6 alleles (BB, green) negatively impact oocyte number. CC: homozygous CAST, BB: homozygous B6, BC: heterozygous CAST/B6 (D). Effect of QTL genotype on oocyte number. The genotype of the QTL region is indicated on the X-axis; oocyte number is on Y-axis. CC: homozygous CAST, BB: homozygous B6, BC: heterozygous CAST/B6 (E).

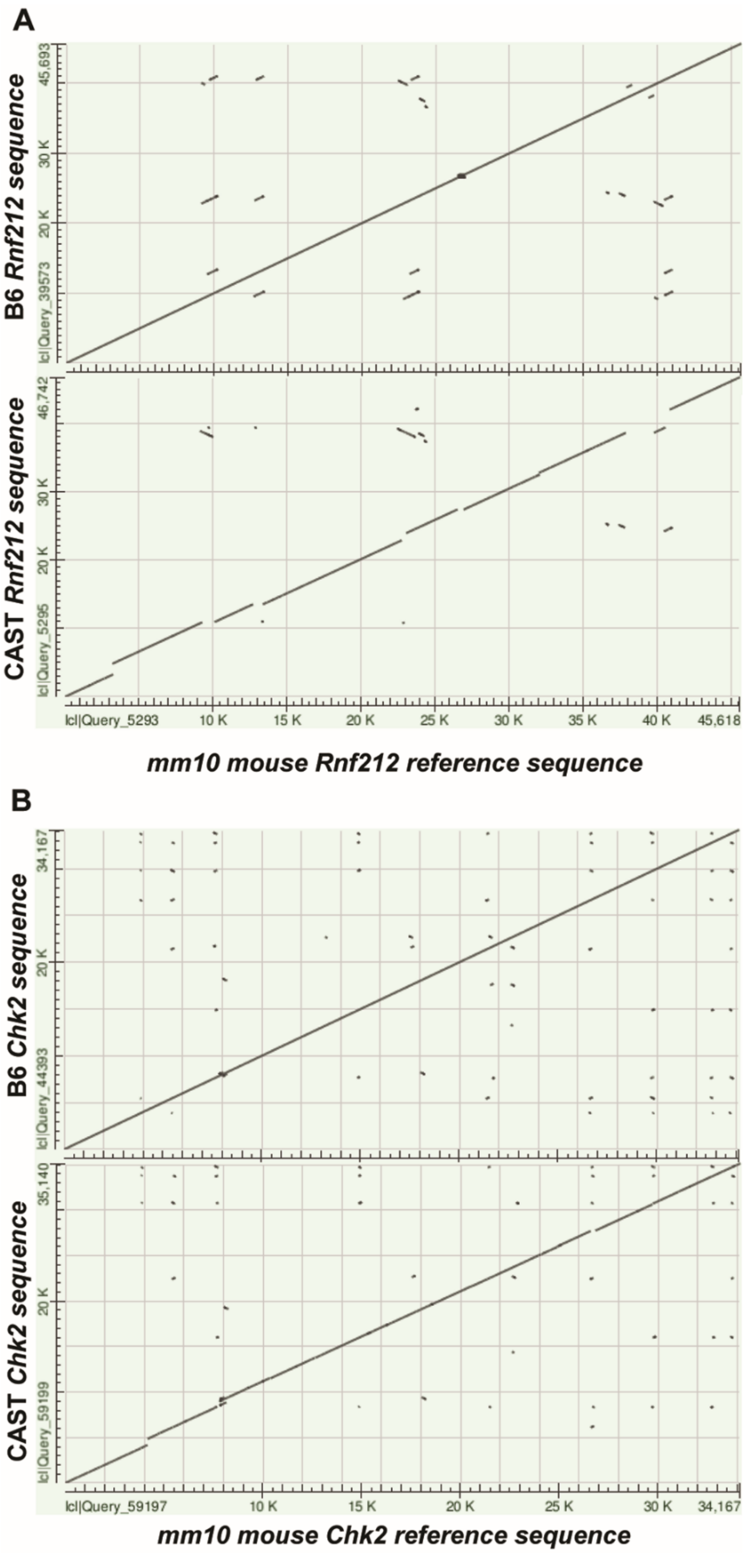


**Fig. S3.** *Rnf212* and *Chk2* genomic sequences are conserved between the B6 and CAST mice. (A-B) Dot Matrix view from Blast 2 Sequences showing the alignment of *Rnf212* (A) and *Chk2* (B) genes in B6 and CAST mice to the mouse reference sequence (mm10).

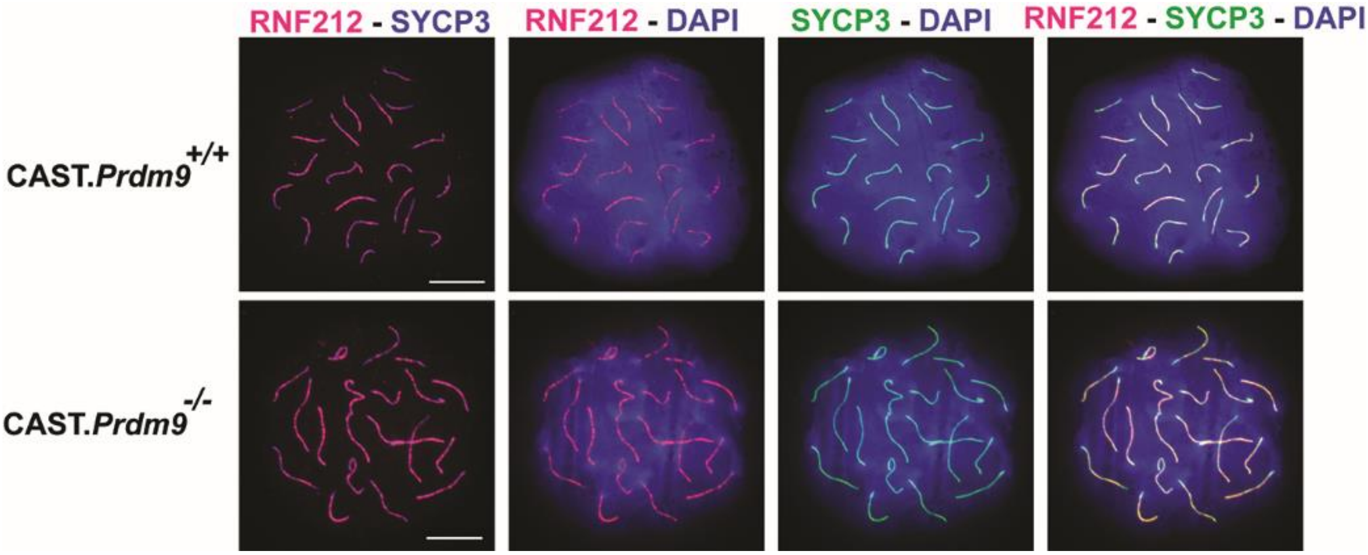


**Fig. S4.** RNF212 protein expression and co-localization patterns are unaffected in CAST.*Prdm9*$^{-/-}$ meiotic oocytes. CAST wild type and *Prdm9*$^{-/-}$ meiotic oocyte chromatin spreads are immunolabeled with antibodies against DSB repair protein RNF212 (red), and SC protein SYCP3 (blue or green). DNA is stained with DAPI (blue). Note co-localization of RNF212 and SYCP3. Scale bars: 10μm.

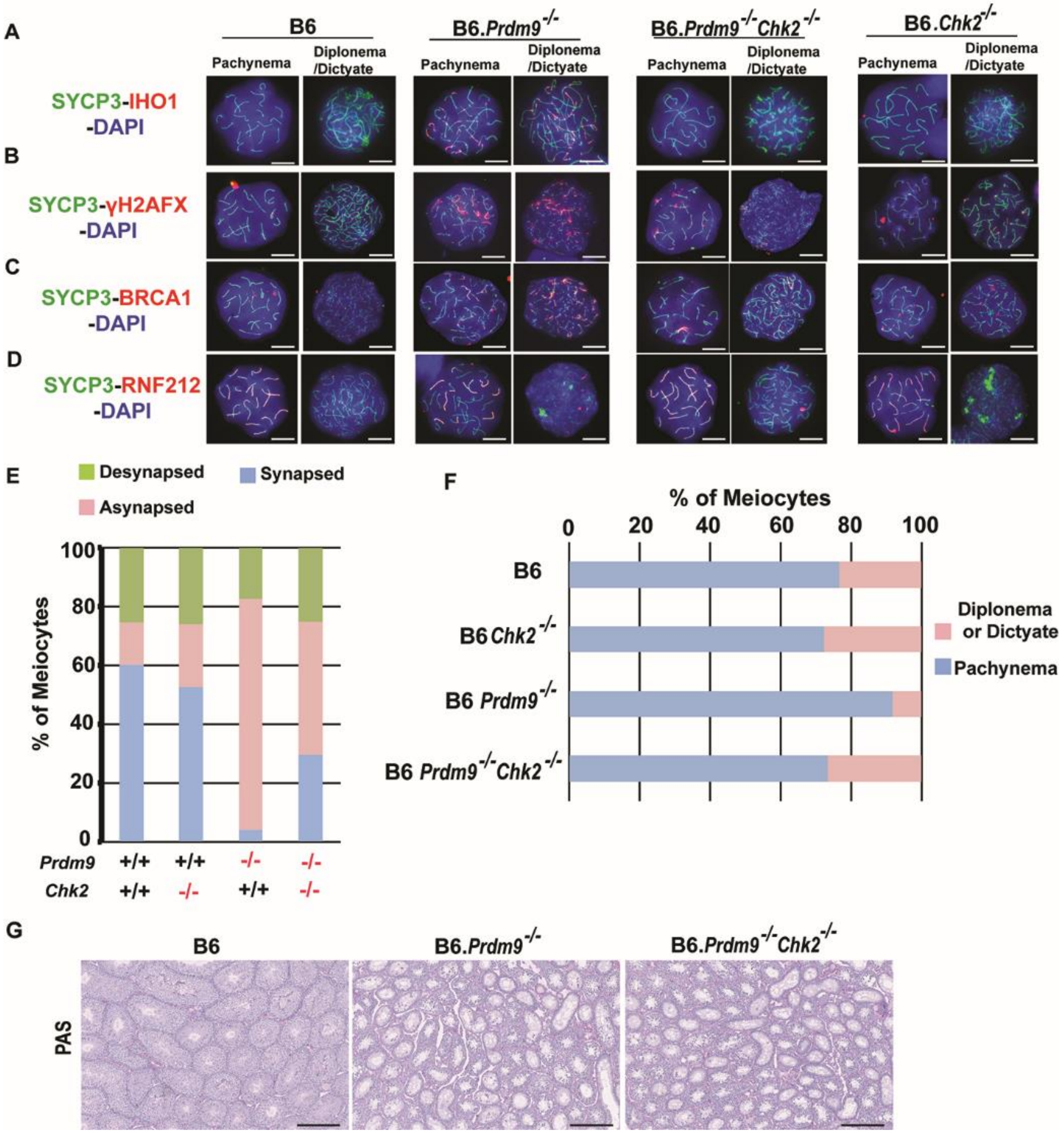

A
B
C
D
B6
B6.Prdm9-/-
B6.Prdm9-/-Chk2-/-
B6.Chk2-/-
Pachynema
Diplonema /Dictyate
SYCP3-IHO1 -DAPI
SYCP3-γH2AFX -DAPI
SYCP3-BRCA1 -DAPI
SYCP3-RNF212 -DAPI
E
Desynapsed
Synapsed
Asynapsed
% of Meiocytes
0
20
40
60
80
100
Prdm9 +/+ +/+ -/- -/-
Chk2 +/+ -/- +/+ -/-
F
% of Meiocytes
0 20 40 60 80 100
B6
B6 Chk2-/-
B6 Prdm9-/-
B6 Prdm9-/-Chk2-/-
Diplonema or Dictyate
Pachynema
G
B6
B6.Prdm9-/-
B6.Prdm9-/-Chk2-/-
PAS

**Fig. S5.** Sexually dimorphic phenotypes in B6.*Prdm9*$^{-/-}$*Chk2*$^{-/-}$ mice. (A) Co-immunolabeling detection of the DSB-associated protein IHO1 (red) and SYCP3 (green) in pachytene and dictyate oocyte chromatin spreads from the indicated wild type and mutant females. DNA is stained with DAPI (blue). (B) Co-immunolabeling detection of the DSB-associated protein γ-H2AFX (red) and SYCP3 (green) in pachytene and dictyate oocyte chromatin spreads from the indicated wild-type and mutant females. DNA is stained with DAPI (blue). (C) Co-immunolabeling detection of DSB repair protein BRCA1 (red) and SYCP3 (green) in pachytene and dictyate oocyte chromatin spreads from the indicated wild-type and mutant females. DNA is stained with DAPI (blue). (D) Co-immunolabeling detection of the crossover-promoting protein RNF212 (red) and SYCP3 (green) in pachytene and dictyate oocyte chromatin spreads from the indicated wild type and mutant females. DNA is stained with DAPI (blue). All scale bars = 10 µm. (E) Stacked bar chart showing frequencies of chromosomal synapsis status in pachytene oocytes in P0 ovary. The genotypes of the mice tested are indicated below the graph. (F) Stacked bar chart showing frequencies of different meiotic oocytes in P0 ovary. The genotypes of the mice tested are indicated on the left side of the graph. (G) PAS-stained histological sections of mouse testis sections of the indicated genotypes at 8 weeks of age. Scale bars = 100 µm. All mutants have meiotic arrest at testis epithelial Stage IV.

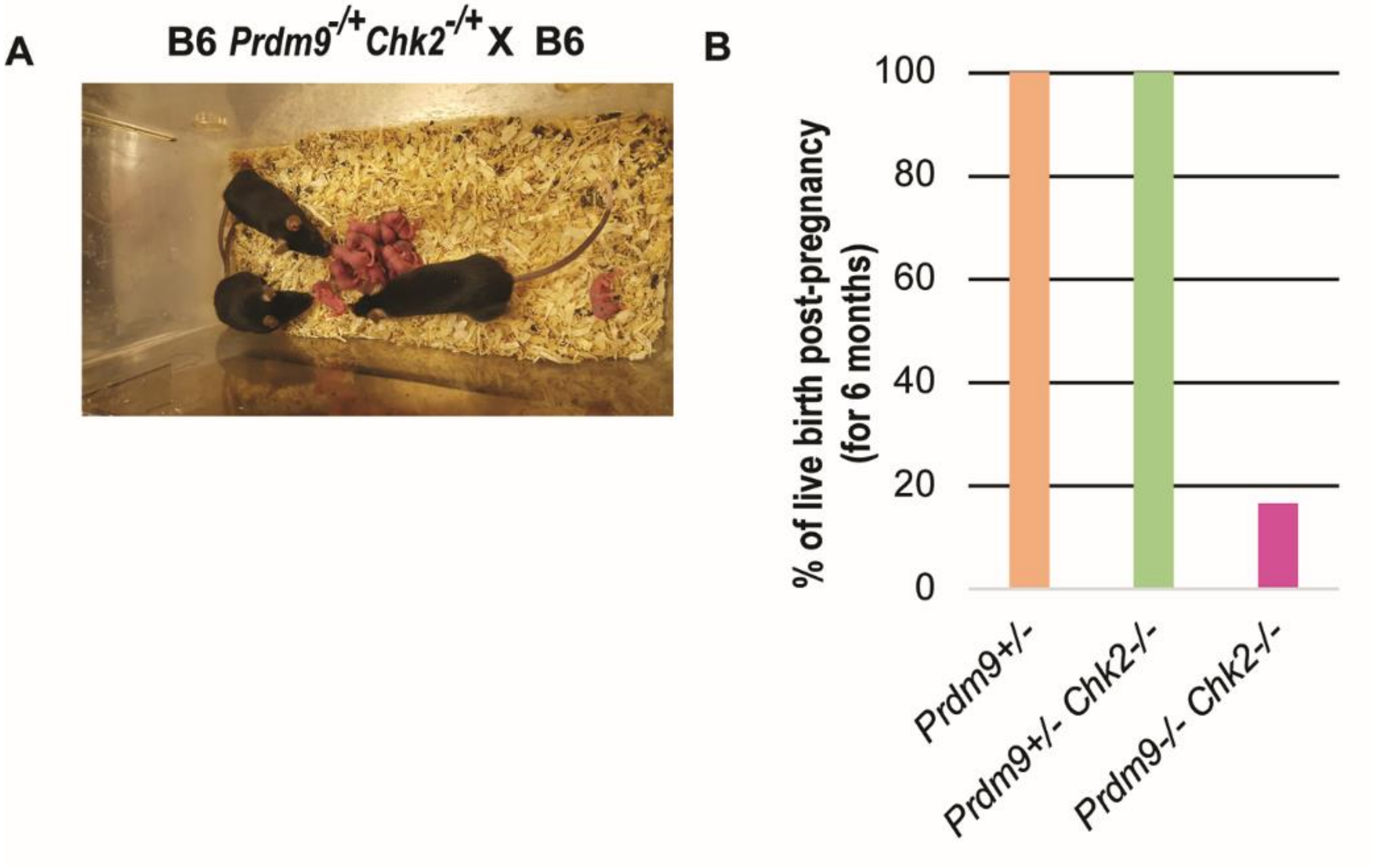


**Fig. S6.** Viability and fertility of embryos produced by B6.*Prdm9*$^{-/-}$*Chk2*$^{-/-}$ female mice. (A) Pups produced by the female B6. *Prdm9*$^{+/-}$ *Chk2*$^{+/-}$ offspring of B6. *Prdm9*$^{-/-}$ *Chk2*$^{-/-}$ female mice. (B) Bar graph showing percentage of live births noted after observed pregnancy (at 16 dpc) in female mice mated to wild type males. The data presented here represent observations over 6 months. The genotypes of mice tested are indicated below the graphs.

| Genotype | Hotspot Peak Number | % B6 Peak Number | Fold Reduction in Peak Number |
|---|---|---|---|
| B6 | 18670 | 100 | - |
| *Prdm9*$^{EP/EP}$ | 2591 | 13.9 | 7.21 |

**Table S1.** Hotspot peak number in wild-type and mutant spermatocytes
Out of 18,838 known PRDM9-dependent H3K4me3 peaks, the table above shows the number of detectable H3K4me3 ChIP-seq peaks in wild-type B6, and *Prdm9*$^{EP/EP}$ spermatocytes isolated from 14dpp males. Peaks were called by MACS 1.4.

| Genotype | Hotspot Peak Number | Avg. tags per peak (RPM) | Fold Reduction avg. tags per peak (RPM) |
|---|---|---|---|
| B6, EP hotspots | 2591 | 5.08 | - |
| *Prdm9*$^{EP/EP}$, EP hotspots | 2591 | 0.825 | 6.16 |

**Table S2**. Hotspot peak intensity reduction in mutant spermatocytes
Hotspot peak intensity (average tags per peak) at detectable PRDM9-dependent H3K4me3 peaks in *Prdm9*$^{EP/EP}$ spermatocytes, relative to average peak intensity at those peaks in B6.

| Genetic Background | Duration (months) | Females in cross | Total Pups | Total Litters | Avg. pups/litter | OFM | LFM | Pups lost |
|---|---|---|---|---|---|---|---|---|
| B6 | 6 | 1 | 5 | 1 | 5 | 0.83 | 0.17 | 0 |
| B6 | 6 | 1 | 28 | 5 | 5.6 | 4.67 | 0.83 | 5 |
| B6 | 6 | 1 | 10 | 2 | 5 | 1.67 | 0.33 | 0 |
| B6 | 6 | 1 | 10 | 2 | 5 | 1.67 | 0.33 | 2 |
| B6 | 6 | 1 | 24 | 5 | 4.8 | 4.00 | 0.83 | 9 |
| B6 | 6 | 1 | 20 | 5 | 4 | 3.33 | 0.83 | 2 |
| **AVG** | | | | | | **2.69** | **0.56** | **3** |

| Genetic Background | Duration (months) | Females in cross | Total Pups | Total Litters | Avg. pups/litter | OFM | LFM | Pups lost |
|---|---|---|---|---|---|---|---|---|
| B6xWSB-F2 | 6 | 1 | 24 | 6 | 4 | 4.00 | 1.00 | 0 |
| B6xWSB-F2 | 6 | 2 | 41 | 10 | 4.1 | 3.42 | 0.83 | 0 |
| B6xWSB-F2 | 6 | 2 | 74 | 13 | 5.7 | 6.17 | 1.08 | 3 |
| B6xWSB-F2 | 6 | 2 | 41 | 8 | 5.1 | 3.42 | 0.67 | 1 |
| B6xWSB-F2 | 6 | 1 | 28 | 6 | 4.7 | 4.67 | 1.00 | 1 |
| B6xWSB-F2 | 7+ | 2 | 79 | 9 | 8.8 | 5.64 | 0.64 | 5 |
| **AVG** | | | | | | **4.55** | **0.87** | **1.67** |

**Table S3**. *Prdm9*$^{EP/EP}$ female fertility tests; B6 and B6xWSB-F2 genetic backgrounds
Females were crossed at 5 weeks of age with a sexually mature (>8 weeks), wild-type B6 male, in pairs or trios as noted. Crosses were allowed to continue for at least six months, and longer if necessary until the females stopped producing offspring. The number of pups in each cross that died before wean are noted in the last column; all other offspring appeared grossly normal and healthy at wean. All offspring in the B6 background, and the offspring from at least 3 litters per cross in the B6xWSB-F2 background, were genotyped for *Prdm9*$^{EP}$ to confirm maternal homozygosity. As expected, all genotyped offspring were heterozygous for *Prdm9*$^{EP}$.
OFM: offspring per female per month; LFM: litters per female per month.

| Genotype | Total female | Average pups/litter | Standard Deviation | Total litter (N) |
|---|---|---|---|---|
| B6 | 3 | 7.5 | 1.1 | 15 |
| B6.*Prdm9*EP/EP | 6 | 4.6 | 2 | 20 |
| B6WSBF2.*Prdm9*EP/EP | 10 | 5.5 | 3.2 | 53 |
| B6.*Prdm9-/-* | 6 | 0 | 0 | 0 |
| C3H | 3 | 8.9 | 1.3 | 15 |
| C3H.*Prdm9-/-* | 7 | 0 | 0 | 0 |
| CAST | 3 | 5.9 | 1.4 | 15 |
| CAST.*Prdm9-/-* | 4 | 6.4 | 1.2 | 7 |
| B6C3F1.*Prdm9-/-* | 3 | 0 | 0 | 0 |
| B6CASTF1.*Prdm9-/-* | 3 | 7.5 | 2.1 | 7 |
| C3HCASTF1.*Prdm9-/-* | 2 | 8 | 1.3 | 6 |
| B6.*Chk2-/-* | 3 | 8 | 1.4 | 5 |
| B6.*Prdm9-/-Chk2-/-* | 10 | 1.8 | 0.8 | 10 |

**Table S4**. Fertility data for wild-type and mutant females in this study
Litter size and variation, and total number of litters, for females of all genotypes reported in this study.

| MLH1 | B6 | B6.*Prdm9*-/- | B6 *Prdm9* EP/EP |
|---|---|---|---|
| Average | 27,6 | 16,2 | 30,5 |
| standard deviation | 3,8 | 8,6 | 4,6 |
| Total meiocytes (n) | 42 | 49 | 30 |
| Total females (N) | 2 | 3 | 2 |

| MLH1 | CAST | CAST.*Prdm9*-/- | B6.*Chk2*-/- | B6.*Prdm9*-/- *Chk2*-/- |
|---|---|---|---|---|
| Average | 29,3 | 27,2 | 26,8 | 21,9 |
| standard deviation | 3,1 | 3,3 | 3,5 | 8,7 |
| Total meiocytes (n) | 32 | 32 | 21 | 41 |
| Total females (N) | 2 | 2 | 2 | 3 |

| MLH3 | B6 | B6.*Prdm9*-/- | B6.*Chk2*-/- | B6.*Prdm9*-/- *Chk2*-/- |
|---|---|---|---|---|
| Average | 33,3 | 13,6 | 32,3 | 23,8 |
| standard deviation | 4,1 | 7,5 | 5,5 | 6,7 |
| Total meiocytes (n) | 30 | 46 | 22 | 50 |
| Total females (N) | 3 | 3 | 2 | 3 |

**Table S5.** Average Number of MLH1 and MLH3 Foci for wild-type and mutant females in this study
n: represents total count of meiocytes per genotype, while N: represents total number of animals assayed per genotype in this study.

| Genotype | Weight of 2 Adult Testes (mg) |
|---|---|
| B6WSBF2.*Prdm9*$^{EP/EP}$ | 61.6 |
| B6WSBF2.*Prdm9*$^{EP/EP}$ | 45.2 |
| B6WSBF2.*Prdm9*$^{EP/EP}$ | 85.2 |
| B6WSBF2.*Prdm9*$^{EP/EP}$ | 65.2 |
| B6WSBF2.*Prdm9*$^{EP/EP}$ | 54.4 |
| B6WSBF2.*Prdm9*$^{EP/EP}$ | 69.7 |
| B6WSBF2.*Prdm9*$^{EP/EP}$ | 40.6 |
| B6WSBF2.*Prdm9*$^{EP/EP}$ | 52.6 |
| B6WSBF2.*Prdm9*$^{+/EP}$ | 255 |
| B6WSBF2.*Prdm9*$^{+/EP}$ | 233 |
| B6WSBF2.*Prdm9*$^{+/+}$ | 297 |
| B6WSBF2.*Prdm9*$^{+/+}$ | 264 |

**Table S6.** Testis weights in B6WSBF2 males. Weights of combined adult (>8wk) testes are shown for eight B6WSBF2.*Prdm9*$^{EP/EP}$ males, as well as two each of heterozygous and wild-type males for comparison.
